# Reproducibility of 4D Flow MRI-based Personalized Cardiovascular Models; Inter-sequence, Intra-observer, and Inter-observer variability

**DOI:** 10.1101/2024.06.13.597551

**Authors:** Belén Casas, Kajsa Tunedal, Federica Viola, Gunnar Cedersund, Carl-Johan Carlhäll, Matts Karlsson, Tino Ebbers

**Affiliations:** Department of Health, Medicine and Caring Sciences, Linköping University, Linköping, Sweden; Center for Medical Image Science and Visualization (CMIV), Linköping University, Linköping, Sweden; Department of Biomedical Engineering, Linköping University, Linköping, Sweden; School of Medical Sciences and Inflammatory Response and Infection Susceptibility Centre (iRiSC), Faculty of Medicine and Health, Örebro University, Örebro, Sweden; Department of Clinical Physiology in Linköping, and Department of Health, Medicine and Caring Sciences, Linköping University, Linköping, Sweden; Department of Management and Engineering, Linköping University, Linköping, Sweden

## Abstract

Subject-specific parameters in lumped hemodynamic models of the cardiovascular system can be estimated using data from experimental measurements, but the parameter estimation may be hampered by the variability in the input data. In this study, we investigate the influence of inter-sequence, intra-observer, and inter-observer variability in input parameters on estimation of subject-specific model parameters using a previously developed approach for model-based analysis of data from 4D Flow MRI acquisitions and cuff pressure measurements. The parameters describe left ventricular time-varying elastance and aortic compliance. Parameter reproducibility with respect to variability in the MRI input measurements was assessed in a group of ten healthy subjects. The subject-specific parameters had coefficient of variations between 2.5% and 34.9% in the intra- and inter-observer analysis. In comparing parameters estimated using data from the two MRI sequences, the coefficients of variation ranged between 3.6% and 41%. The diastolic time constant of the left ventricle and the compliance of the ascending aorta were the parameters with the lowest and the highest variability, respectively. In conclusion, the modeling approach allows for estimating left ventricular elastance parameters and aortic compliance from non-invasive measurements with good to moderate reproducibility concerning intra-user, inter-user, and inter-sequence variability.

## INTRODUCTION

Computational modeling of the cardiovascular system has gained increased relevance as a tool to understand physiological and pathophysiological changes, providing new insights into diagnosing and treating cardiovascular diseases. Patient-specific representations are being increasingly used, with model personalization being driven by individually acquired measurements that can be obtained invasively or non-invasively ^1^. Among current cardiovascular modeling approaches, lumped parameter models have been proposed as a fast, relatively simple method to simulate global hemodynamics while keeping computational demands low ^2^. Several lumped parameter modeling approaches have been developed for studying global hemodynamics. These generally combine lumped parameter models of the cardiovascular system with subject-specific input measurements derived from ultrasound measurements, cardiac magnetic resonance (CMR) acquisitions and invasive pressure catheter measurements in the heart or the aorta ^3–6^.

Following personalization, the subject-specific model parameters can be used as cardiovascular biomarkers with potential application for diagnostic and prognostic purposes ^7–12^. However, in order for these biomarkers to be reliable and successfully applied in healthcare, the methods for identifying them should be robust against errors and variability in these input data ^13^. Previous studies have identified several sources of variability in computational models and highlighted the importance of considering errors in input measurements, also referred to as observational uncertainty, as a source of uncertainty in model-derived parameters ^14–17^. Observational uncertainty is of interest in the evaluation of parameters derived from lumped parameter cardiovascular models, as personalization typically relies on a variety of anatomical and functional measurements that are susceptible to acquisition and analysis errors.

In a previous study, we presented an approach to personalize a lumped parameter model of the systemic circulation based on non-invasive measurements, which comprise CMR morphological images and flow data from four-dimensional magnetic resonance imaging (4D Flow MRI) ^6^. The given approach allows for the identification of several cardiovascular parameters which are relevant for diagnostic purposes, such as parameters describing the contractile properties of the left ventricle in terms of its time-varying elastance ^18^, and the compliance of the ascending aorta.

As a continuation of this study, we aimed to assess the reproducibility of the estimated parameters in relation to variability in the analysis and acquisition settings of the input images to the model. For this purpose, the parameters were investigated in a study of ten healthy volunteers examined with two 4D Flow MRI sequences with different scan parameters, which were then analyzed by two different observers in order to evaluate intra-observer, inter-observer and inter-sequence variability.

## RESULTS

### Variability in input image-derived measures

The results of the intra-, inter-observer variability, and repeatability using the spoiled gradient echo (SGRE) and echo-planar imaging (EPI) 4D flow MRI sequences for the input measures to the model are shown in Table 1. None of the input parameters showed a significant mean difference in these investigations (P > 0.3 in all cases). The input parameter with the lowest coefficient of variation (CoV) in the intra-observer analysis was the effective orifice area of the aortic valve (EOA) (1.5%), while the lowest coefficient of variation in the inter-observer analysis corresponded to end systolic volume (ESV) (3.8%). In both the intra- and inter-observer analysis, as well as when comparing the SGRE and EPI sequences, the LV outflow tract area (A_LVOT_) had the highest coefficient of variation (2.8, 11.3 and 11.6%) among all the input measures. For all input parameters, results of CoV from the inter-observer study and the comparison between the SGRE and EPI sequences were higher than their corresponding values from the intra-observer analysis. The variations in the length of the cardiac cycle between the acquisitions using the SGRE and EPI sequences had a CoV of 3.7% across all subjects, indicating a low variability in heart rate between the scans.

**Table 1:** Repeatability of input measures to the model derived from the MRI acquisitions comparing intra-, inter-observer analysis and the SGRE and EPI sequences. A_LVOT_: left ventricular outflow tract area, ESV: left ventricular end systolic volume, EOA: effective orifice area of the aortic valve, T: length of cardiac cycle.

| Input image-derived measure | Intra-user variability |  |  |
| --- | --- | --- | --- |
|  | Mean difference (95% confidence interval) | P value | CoV (%) |
| $A_{LVOT}$ (cm<sup>2</sup>) | -0.1 (-1.2 to 1) | 0.8 | 2.8 |
| ESV (mL) | -0.008 (-15.9 to 16) | 0.9 | 3 |
| EOA (cm<sup>2</sup>) | 0.04 (-0.5 to 0.5) | 0.9 | 1.5 |
|  | Inter-user variability |  |  |
|  | Mean difference (95% confidence interval) | P value | CoV (%) |
| $A_{LVOT}$ (cm<sup>2</sup>) | -0.06 (-1.4 to 1.3) | 0.9 | 11.3 |
| ESV (mL) | 0.8 (-15.4 to 17) | 0.9 | 3.8 |
| EOA (cm<sup>2</sup>) | -0.3 (-0.9 to 0.3) | 0.3 | 8.9 |
|  | SGRE-EPI |  |  |
|  | Mean difference (95% confidence interval) | P value | CoV (%) |
| $A_{LVOT}$ (cm<sup>2</sup>) | -0.1 (-1.6 to 1.4) | 0.9 | 11.6 |
| EOA (cm<sup>2</sup>) | 0.2 (-0.5 to 0.9) | 0.6 | 8.6 |
| T (s) | -0.004 (-0.1 to 0.1) | 0.9 | 3.7 |

Fig. 1 illustrates the variability in the volumetric flow waveforms used as input to the model for the intra- and inter-observer analysis and the SGRE-EPI sequence variability. Comparison of the root mean squared error (RMSE) for the input flow waveforms at the mitral valve, the aortic valve and the ascending aorta across all subjects revealed that, on average, the RMSE at the locations of interest was highest between the flow waveforms extracted from the SGRE and the EPI acquisition data, in comparison with the RMSE values found in the intra- and inter-observer analysis. Similarly, RMSE values in the input flow waveforms as a result of inter-observer variability were higher than those due to intra-user variability. The volumetric flows depicted in Fig. 1A, B, C exemplify the higher variability in the input flow waveforms resulting from the inter-observer analysis, as well as from comparison of SGRE and EPI sequences for one subject. In computing net flow volumes, the coefficients of variation for the total mitral flow were 4.1% and 8.9% for the intra- and inter-observer variability, respectively. For the aortic valve, these coefficients of variation were 1.2% and 2.8%, and for the ascending aorta 2.1% and 2.8%.

**Figure 1:**
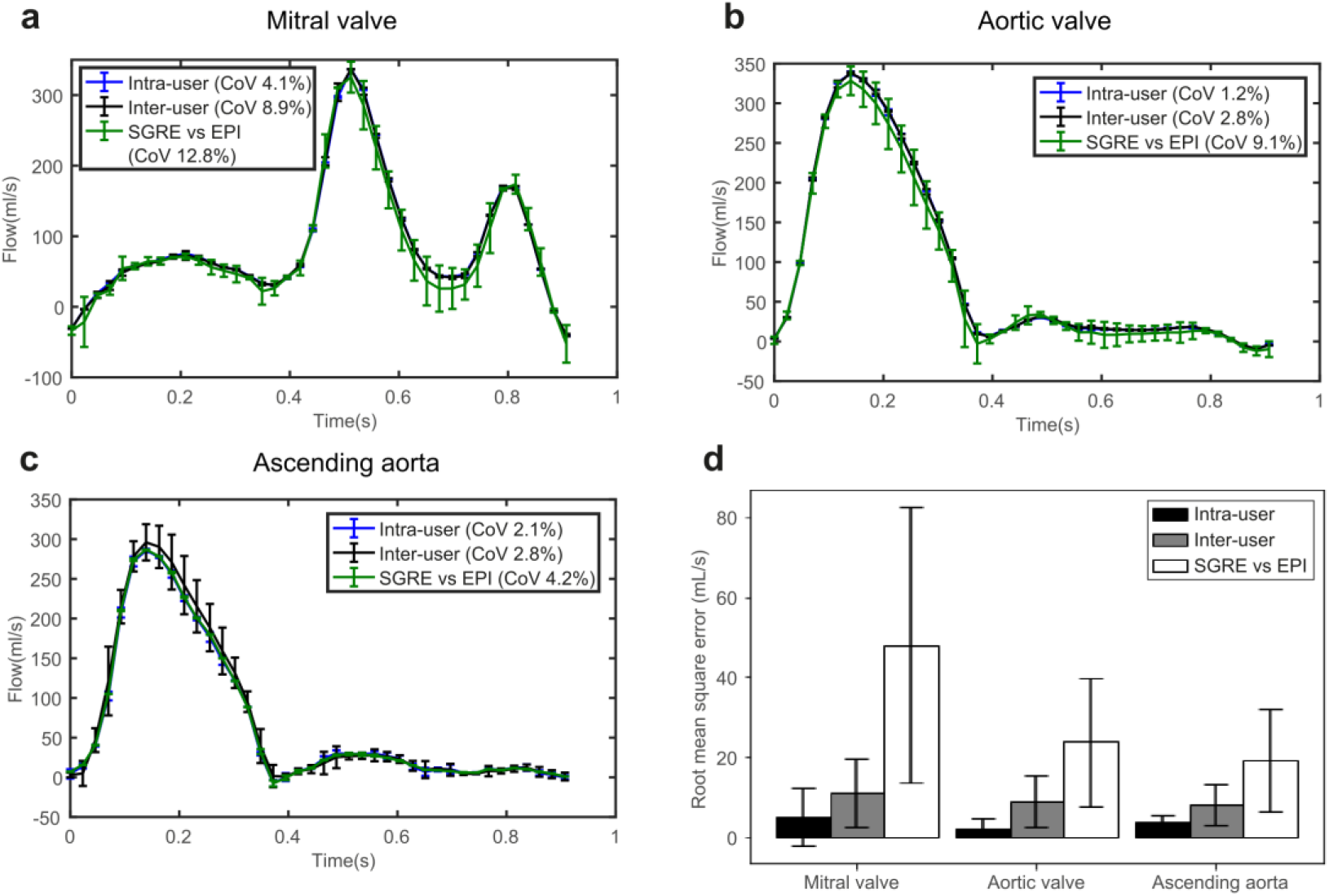
Variability in the volumetric flow waveforms used as input to the model in the mitral valve (A), aortic valve (B) and ascending aorta (C) of a representative subject. The curves and the error bars represent the mean and the standard deviation (±SD) of the volumetric flows for one subject and two different analyses corresponding to the intra-user variability study (blue), inter-user variability study (black) and SGRE-EPI comparison (green). The legend shows the coefficient of variation (Cov) of the net flow volumes, calculated from all 10 subjects. (D) shows the root mean square error (RMSE) of the volumetric flow waveforms across the ten study subjects at the selected measurement sites for these variability studies.

Comparison of the net flow volumes calculated from the SGRE and EPI data yielded coefficients of variations of 12.8%, 9.1% and 4.2% for the mitral valve, the aortic valve and the ascending aorta, respectively.

### Intra- and inter-observer variability in model parameters

Bland-Altman plots for the model-derived parameters obtained with the input data from the intra- and inter-observer study are shown in Fig. 2 and Fig. 3, respectively. Supplementary Table 1 summarizes the bias and limits of agreement obtained from the Bland-Altman analysis. The mean difference values and the coefficients of variation are reported in Table 2. No significant differences were found between the mean values of estimated the parameters, computed over the analyzed subjects, in either the intra- or the inter-observer analysis (P>0.4 in all cases). The parameter with the lowest coefficient of variation was the ventricular diastolic time constant (2.5% and 3.9% for the intra- and inter-observer analysis, respectively), while the parameter with the highest coefficient of variation when analysing both the intra- and inter-observer variability was the compliance of the ascending aorta (21.8% and 34.9%, respectively). The CoV in the rest of the parameters varied between 4.7% and 13.7% for the intra-observer analysis and 5.3% and 19.4% for the inter-observer analysis. For every parameter, results of the CoV from the inter-observer variability analysis were slightly higher than those from the corresponding inter-observer study. Bland-Altman intra-observer comparisons of the model parameters revealed a good agreement, with a bias lower than 2.7% for all parameters. For the inter-observer study, the bias was lower than 10.6% for all the estimated model parameters.

**Figure 2:**
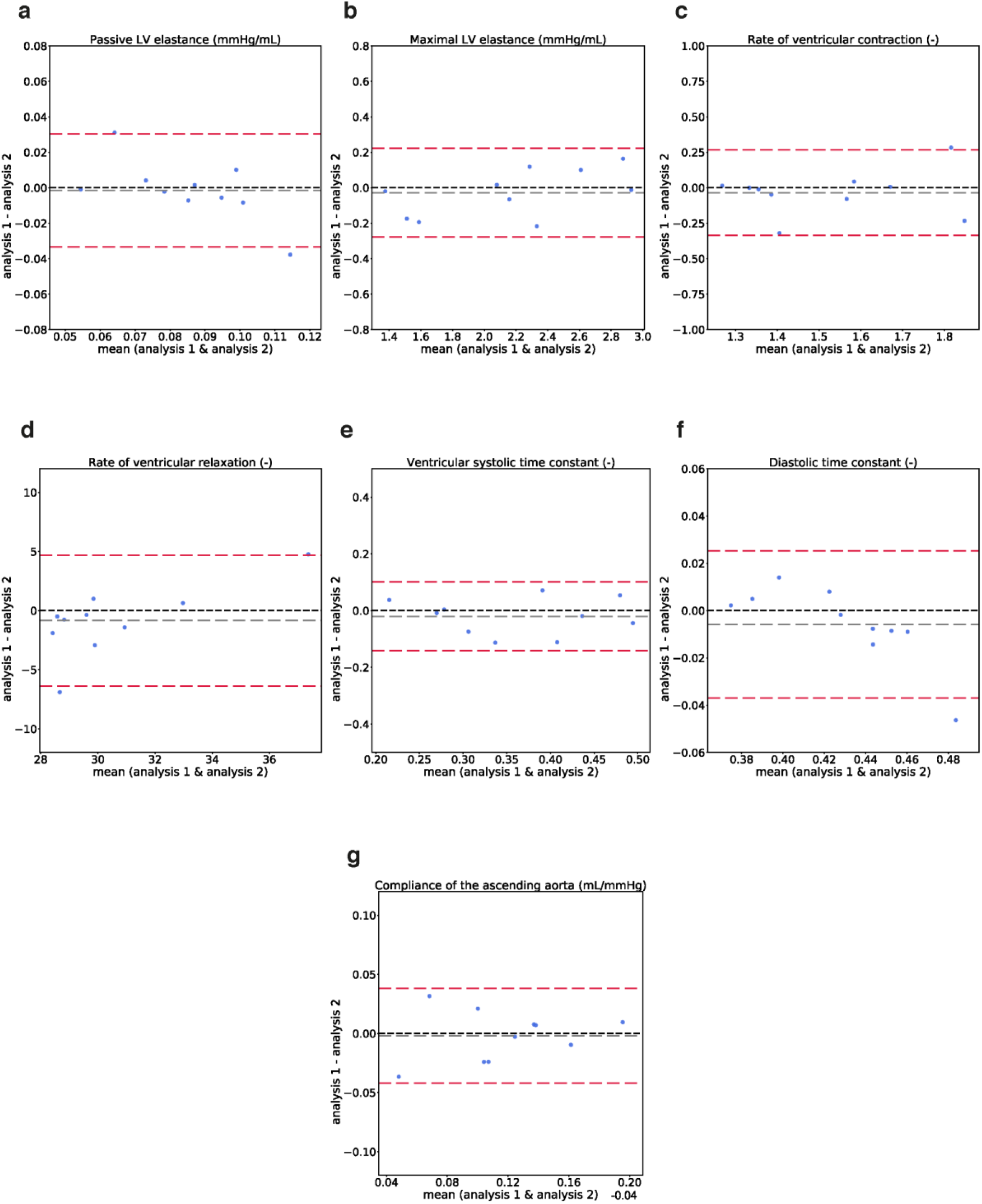
Bland-Altman plots showing agreement in the estimated model parameters using model inputs from the intra-observer variability study. Grey dotted lines represent bias (*d̅*) and red dotted lines represent limits of agreement (*d̅* ± 1.96*SD*). LV: left ventricle.

**Figure 3:**
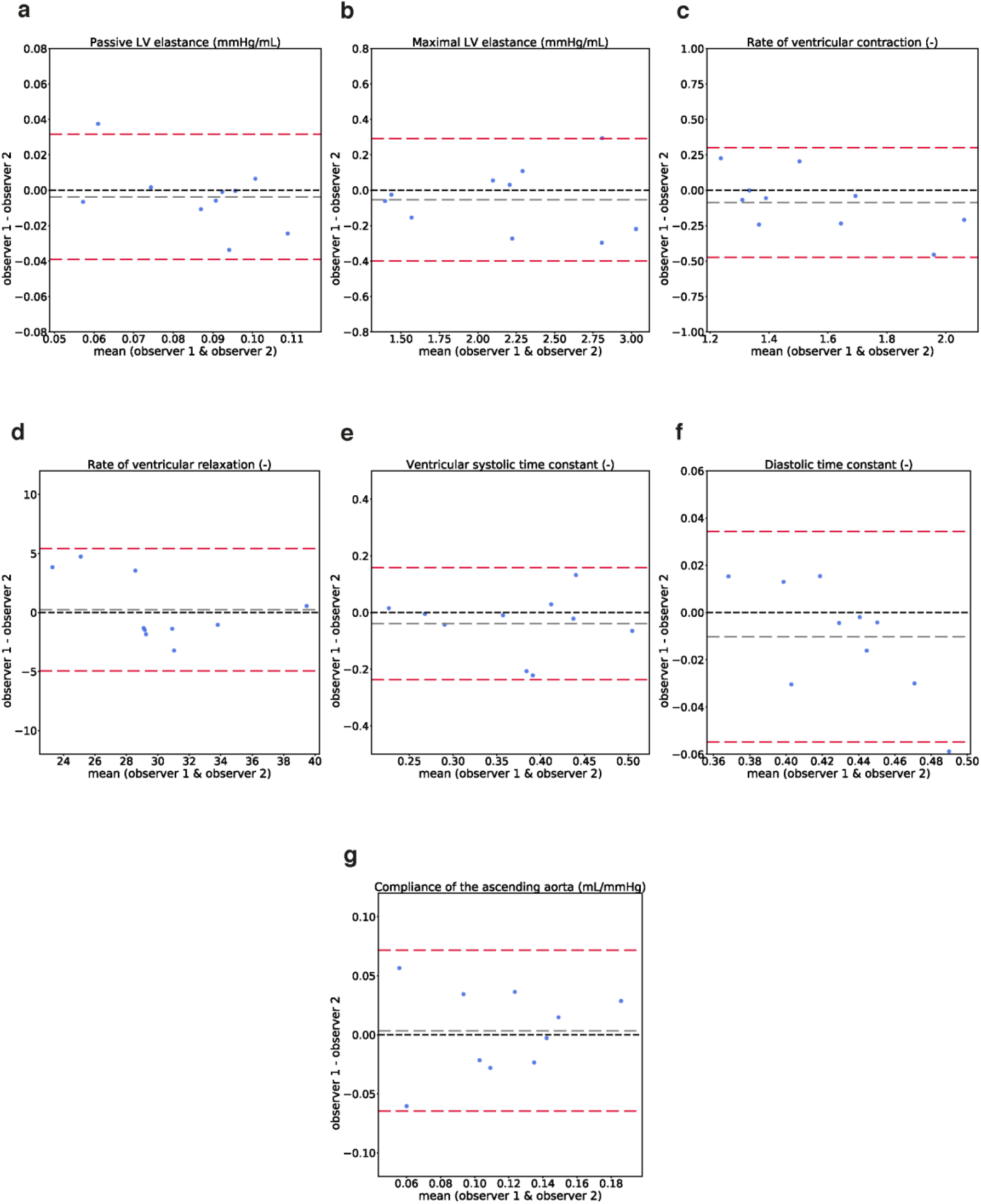
Bland-Altman plots showing agreement in the estimated model parameters using model inputs from the inter-observer variability study. Grey dotted lines represent bias (*d̅*) and red dotted lines represent limits of agreement (*d̅* ± 1.96*SD*). LV: left ventricle.

**Table 2:** Repeatability of model-derived parameter estimations comparing intra- and inter-observer analysis. CoV: coefficient of variation, LV: left ventricle.

| Model<br>parameter | Intra-observer |  |  | Inter-observer |  |  |
| --- | --- | --- | --- | --- | --- | --- |
|  | Mean difference<br>(95% confidence<br>interval) | P value | CoV<br>(%) | Mean difference<br>(95% confidence<br>interval) | P value | CoV<br>(%) |
| Passive LV<br>elastance<br>(mmHg/mL) | -0.007<br>(-0.03 to 0.01) | 0.4 | 13.7 | -0.004<br>(-0.02 to 0.01) | 0.7 | 17.2 |
| Maximal LV<br>elastance<br>(mmHg/mL) | -0.03<br>(-0.5 to 0.5) | 0.9 | 4.7 | -0.05<br>(-0.6 to 0.5) | 0.8 | 5.3 |
| Rate of<br>ventricular<br>contraction (-) | -0.03<br>(-0.2 to 0.2) | 0.7 | 6.9 | -0.09<br>(-0.4 to 0.2) | 0.5 | 9.3 |
| Rate of<br>ventricular<br>relaxation (-) | -0.8<br>(-3.8 to 2.1) | 0.6 | 6.8 | 0.2<br>(-4.1 to 4.6) | 0.9 | 7.1 |
| Ventricular<br>systolic time<br>constant | -0.02<br>(-0.1 to 0.07) | 0.6 | 12.9 | -0.04<br>(-0.1 to 0.06) | 0.4 | 19.4 |
| Ventricular<br>diastolic time<br>constant (-) | -0.006<br>(-0.04 to 0.03) |  | 2.5 | -0.01<br>(-0.05 to 0.02) | 0.5 | 3.9 |
| Compliance of<br>the ascending<br>aorta<br>(mL/mmHg) | -0.002<br>(-0.04 to 0.04) |  | 21.8 | 0.003<br>(-0.04 to 0.04) | 0.9 | 34.9 |

### Inter-sequence variability in model parameters

Bland-Altman comparisons of the model parameters estimated using input data from the SGRE and EPI sequences are shown in Fig. 4. The bias and limits of agreement calculated from this analysis are reported in Supplementary Table 2. Table 3 summarizes the mean differences and CoV values of the estimated parameters. No significant differences were found between the values of the parameters estimated using input data from the two different sequences (P>0.2 in all cases). The ventricular diastolic time constant had the lowest coefficient of variation (3.6%), while the most variable parameter was the compliance of the ascending aorta (CoV 41%). For the rest of the model parameters, the coefficients of variance were in the range 7.5-14.9%.

**Figure 4:**
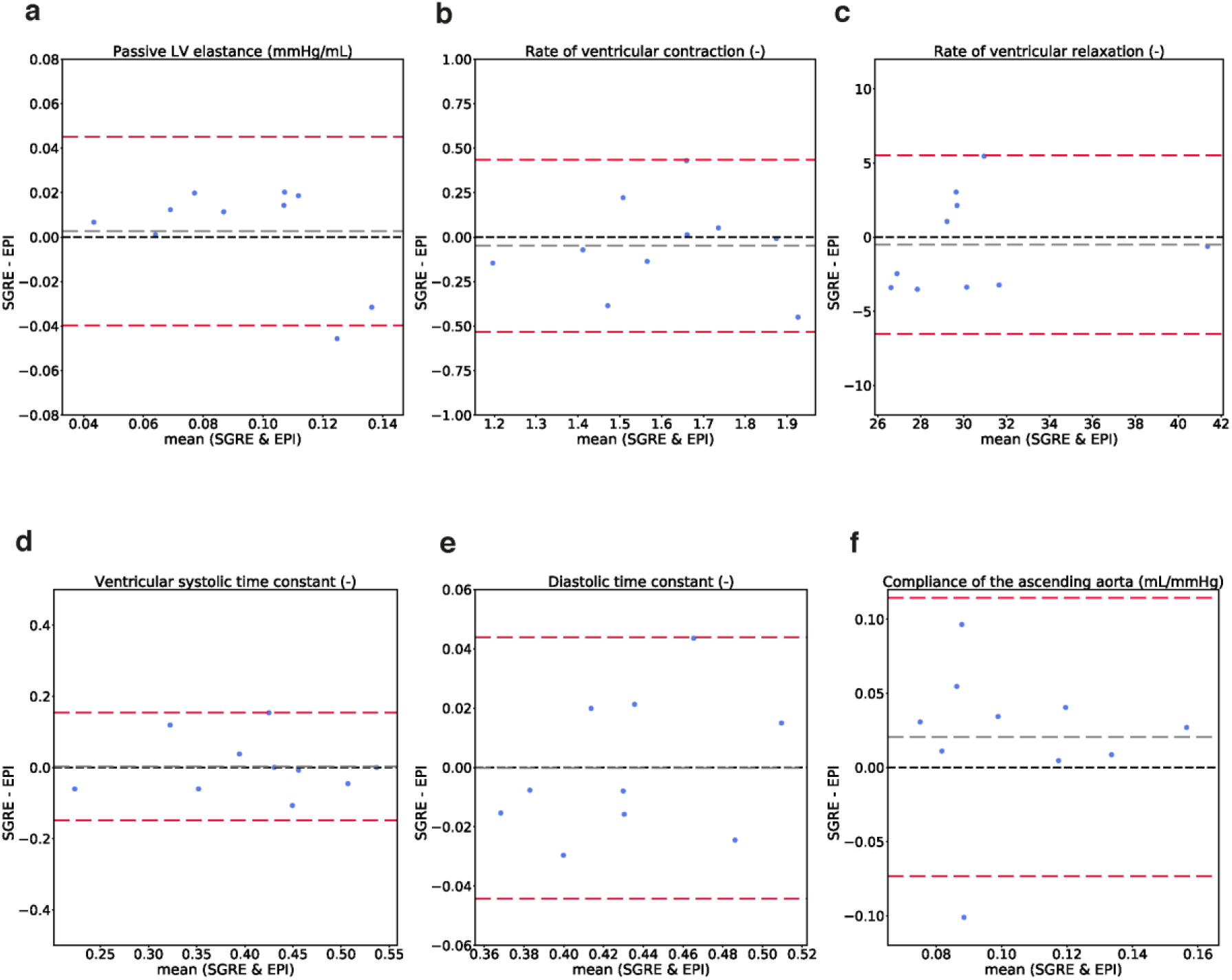
Bland-Altman plots showing agreement in the estimated model parameters using model inputs derived from the SGRE and the EPI measurements. Grey dotted lines represent bias (*d̅*) and red dotted lines represent limits of agreement (*d̅* ± 1.96*SD*). LV: left ventricle.

**Table 3:** Repeatability of model-derived parameter estimations comparing results from SGRE and EPI sequences. CoV: coefficient of variation, LV: left ventricle.

| Model parameter | SGRE-EPI |  |  |
| --- | --- | --- | --- |
|  | Mean difference (95% confidence interval) | P value | CoV (%) |
| Passive LV elastance (mmHg/mL) | 0.003 ( -0.04 to 0.04) | 0.9 | 14.3 |
| Rate of ventricular contraction (-) | -0.05 (-0.3 to 0.2) | 0.7 | 10.9 |
| Rate of ventricular relaxation (-) | -0.5 (-4.7 to 3.7) | 0.8 | 7.5 |
| Ventricular systolic time constant | -0.003 (-0.1 to 0.1) | 0.9 | 14.9 |
| Ventricular diastolic time constant (-) | -0.0001 (-0.04 to 0.04) | 0.9 | 3.6 |
| Compliance of the ascending aorta<br>(mL/mmHg) | -0.02 (-0.01 to 0.05) | 0.2 | 41 |

## DISCUSSION

This study investigated the reproducibility of model-derived parameters describing left ventricular time-varying elastance and aortic compliance obtained using lumped-parameter modeling in combination with CMR morphological images and flow data from 4D Flow MRI ^6^. The reproducibility was assessed with regard to sources of variability in the acquisition and analysis of the input measurements to the model.

Among subject-specific model parameters determining the time-varying elastance of the left ventricle, low coefficients of variation (CoV <10%) considering both intra- and inter-observer variability were found for the maximal LV elastance, the rate of ventricular contraction, the rate of ventricular relaxation and the ventricular diastolic time constant (Table 2). The passive elastance of the left ventricle and the ventricular systolic time constant showed higher variability, with values of the coefficient of variation in the range 12.9-19.4%. The coefficients of variation for all model-derived parameters were higher for the inter-observer study compared to the intra-observer study. These results are consistent with the variability found in the input measurements to the model. The inter-observer variability in these input measurements across the study subjects, in terms of both the coefficient of variation of the input parameters (Table 1) and the root mean square error of the input flow waveforms (Figure 1D), was consistently higher than that from the intra-observer study. The association between higher variability for the inter-observer study in both the input measurements to the model and the estimated model parameters suggests that a major contribution to the variability in model parameters will be from the data analysis steps rather than the estimation approach.

Assessment of the variability in input measurements due to intra- and inter-user analysis (Table 1) involved segmentation of the left ventricle at end systole to compute the ESV, as well as calculation of the EOA at the aortic valve and the area of the LV outflow tract. The variability in ESV agree well with coefficients of variation reported in previous studies assessing intra- and inter-observer variability in LV volumes derived from CMR ^19,20^. The coefficients of variation resulting from the intra- and inter-observer analysis in the EOA estimations are also in good agreement with values reported by others ^21^. Similarly, the coefficients of variation found for the total flow at the mitral and aortic valves (Figure 1) agree with those from previous studies applying retrospective valve tracking that found coefficients of variation between 3% and 7% ^22^.

Comparison of model-derived parameters obtained based on the input data from the SGRE and EPI acquisitions revealed coefficients of variance comparable to those from the intra- and inter-user observer study. In comparing these sequences, the additional uncertainty in measures derived from the morphological cine MR images (i.e. calculation of the ESV) was not considered.

However, the variability due to the input volumetric flow curves was higher compared to the intra- and inter-user observer studies, resulting in higher coefficients of variation for the total flows as well as higher RMSE values at the mitral valve, the aortic valve and the ascending aorta.

Although heart rate was not significantly different between scans, we hypothesize that these changes in heart rate and physiological changes between scans are a major cause of elevated variability in the flow waveforms, and therefore in model-derived parameters. Parameters that are sensitive to changes in heart rate, such as the ventricular contraction and relaxation rates ^23,24^, may have higher variability under these circumstances, especially for large heart rate variations between sequences.

The compliance of the ascending aorta was found to be the most variable parameter in the intra- and inter-observer study and the comparison between the SGRE and EPI acquisition sequences, with coefficients of variation between 21.8% and 41% (Tables 2 and 3). However, the coefficient of variation for the total flow at the ascending aorta was low (2.8% to 4.2%) and similar in magnitude to the coefficients of variation in the mitral valve and the aortic valve (Figure 1).

These findings suggest that the high variability in the aortic compliance is most probably due to the relatively low temporal resolution (30-40 ms) in our study in combination with only using two aortic planes. Previous approaches to calculating pulse wave velocity (PWV) using two-dimensional phase-contrast MRI (2D PC-MRI) in two aortic locations typically require a temporal resolution lower than 11 ms ^25,26^. However, 4D Flow MRI-based PWV estimation has shown to give similar results with 34-40 ms temporal resolution when utilizing all planes along the aorta ^27–29^. Thus, either a higher temporal resolution or utilizing more information from the 4D flow data could improve the estimation of aortic compliance. Another potential reason for the high variability in the aortic compliance parameter is its correlation with other model parameters governing the aortic flow. A change in the aortic compliance parameter could be compensated for by changes in the aortic resistance, inductance, or the timing of the LV contraction, resulting in variability in the aortic compliance.

The findings in this study were based on intra-modality comparisons and did not include comparisons with a gold standard method to evaluate LV time-varying elastance parameters and compliance of the ascending aorta. Future studies should investigate the variability in the model-based elastance parameters in relation to estimates based on invasively measured LV pressures and volumes recorded over a range of loading conditions and use independently measured data to validate the model predictions. Apart from the observational uncertainty evaluated herein in point estimates of model parameter values, each parameter has an uncertainty that additionally includes factors such as model structure, physiological variation during the scan, and non-normal errors such as background phase offset in the 4D flow MRI^30^. This parameter uncertainty can be evaluated with established methods such as profile likelihood^31^. However, this requires quantification of the input data uncertainty of 4D flow MRI and cuff pressure and handling of the non-normal errors which challenges the normality assumption of the profile likelihood method.

Such an analysis is not yet done, and the parameter uncertainty is therefore not included here. Furthermore, only a small number of subjects were included in the study. Additional studies should assess the variability in the model-derived parameters in a larger subject cohort with a wider spectrum of cardiac volumes and flow characteristics as a step toward assessing the robustness of the modeling approach for clinical use.

In conclusion, the majority of parameters derived from the **modeling** approach using CMR morphological images and flow data from 4D Flow MRI showed low variability with analysis and acquisition settings of the input measurements. Improvements in pulse wave velocity calculation and the use of automatic methods for left ventricular segmentation and analysis of the 4D Flow MRI data may reduce the uncertainty in the input measurements and thereby further decrease the variability in the model parameters. Assessment of the variability in these parameters is essential to establish the credibility of model predictions for their potential use in clinical applications.

## METHODS

### Study population

Ten healthy subjects (4 female, mean age 32 ± 6 years) were included in the study. All participants had no history of cardiovascular disease, no medication for cardiovascular disease and a normal physical examination. The study was approved by the Regional Ethical Review Board in Linköping and complies with the Declaration of Helsinki. All subjects provided written informed consent prior to participation in the study.

### Data acquisition

All examinations were performed using a clinical 1.5 T Philips Ingenia scanner (Philips Healthcare, Best, the Netherlands). The imaging protocol included the acquisition of morphological cine balanced steady-state free precession (bSSFP) images and two 4D Flow MRI examinations for each subject. In addition to the acquisition of the MRI datasets, systolic and diastolic blood pressures (SBP and DBP, respectively) were obtained using cuff-based measurements in the brachial artery.

Morphological bSSFP images comprised a stack of short-axis (SA) images, as well as two-, three- and four-chamber long-axis (LAx) images of the left heart. All bSSPF images were acquired at end-expiratory breath-holds with in-plane resolution of 1 × 1 mm^2^ and slice thickness of 8 mm.

To evaluate the effect of scan settings on the model parameters, we performed two 4D flow MRI acquisitions per subject using two different sequences: a spoiled gradient echo (SGRE) sequence with a k-space segmentation factor of 2, and an echo-planar imaging (EPI) sequence with a read-out factor of 3. The acquisitions were done in random order after each other. Both acquisitions were retrospectively cardiac gated and respiratory navigator gated and followed the recommendations from the 4D flow MRI consensus statements^30,32^. Scan parameters for both sequences were: Sagittal-oblique slab covering the whole heart and the thoracic aorta, velocity encoding (VENC) 120 cm/s and flip angle 5°, and parallel imaging (SENSE) speed up factors of 2 in the AP and RL directions. The echo time (TE) was 3 ms for SGRE and 4 ms for EPI, and the repetition time (TR) was 5 ms for SGRE and 7 ms for EPI. The acquired spatial resolution was 2.9 × 2.9 × 2.9 mm^3^ and the temporal resolution was 40 ms and 30 ms for the SGRE and the EPI sequence, respectively. Excluding navigator efficiency, the scan time was approximately 10 min for SGRE and 8 min for EPI. The average navigator gating efficiency was between 60% and 80%. After acquisition the 4D Flow data were retrospectively reconstructed into 40 time frames and corrected for concomitant gradient fields on the scanner. The resulting data were exported to an off-line station for post-processing and analysis. The post-processing was performed using developed software written in Matlab (The Mathworks Inc., Natick, Massachusetts, USA). The data were corrected for background errors using a weighted second-order polynomial fit to static tissue in the thorax ^33^ and phase wraps using a temporal algorithm ^34^.

### Data analysis

Several functional and morphological parameters were extracted from the imaging data, including parameters that characterize the morphology and the function of the left ventricle (LV) and the aortic valve, as well as volumetric flow waveforms from the heart valves and selected sites in the ascending aorta.

The length of the cardiac cycle was calculated as the average length of the cardiac cycle during the 4D Flow MRI acquisition. The end-systolic volume (ESV) was computed using the short-axis stack images, by manual delineation of the endocardial border at the time of end systole. The segmentations were performed using the freely available segmentation software Segment version 1.9 (Medviso, Lund, Sweden) ^35^.

In the model, the aortic valve is described using the pressure gradient formulation introduced by Garcia et al. ^36^. This formulation requires estimation of the effective orifice area (EOA) of the aortic valve and the cross-sectional area of the LV outflow tract (A_LVOT_). The EOA was calculated as follows:

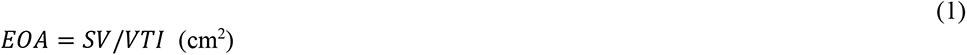

where the stroke volume (SV) was computed as the time integral of the flow at the valve, and VTI was the velocity-time integral of the instantaneous maximal velocity at the valve. A_LVOT_ was calculated as the area of the LV outflow tract (LVOT), measured downstream from the valve during systole ^37^. The contour of the aortic valve was manually delineated and the time integrals of the flow and the velocity, as well as the cross-sectional area of the LVOT, were calculated.

Input flow waveforms to the model include 4D Flow MRI-derived volumetric flow waveforms at the mitral valve, the aortic valve, and the ascending aorta, upstream from the brachiocephalic trunk. Flows through the heart valves were computed using a semi-automatic, retrospective valve tracking approach with correction for through-plane motion. Using this approach, two input points that identify the valve annulus are manually placed in the three-chamber view at the time of end diastole. By automatic tracking of the valve annulus along the successive time frames, in combination with manual segmentation of the valve orifice for each time frame, the approach yields time-resolved curves of the flow through the valve. These calculations were performed using in-house software written in Matlab (The Mathworks Inc., Natick, Massachusetts, USA).

To extract the flow waveforms in the ascending aorta, a 4D PC-MR angiogram was used for anatomical orientation in 3D using commercially available visualization software (EnSight, CEI Inc., NC, USA) to locate the position of the analysis plane. The volumetric flow curves were calculated by manual segmentation of the aortic contour for every cardiac time frame, as described elsewhere ^6^. A schematic summary of the steps involved in the analysis process and the modeling approach is provided in Fig. 5.

**Figure 5:**
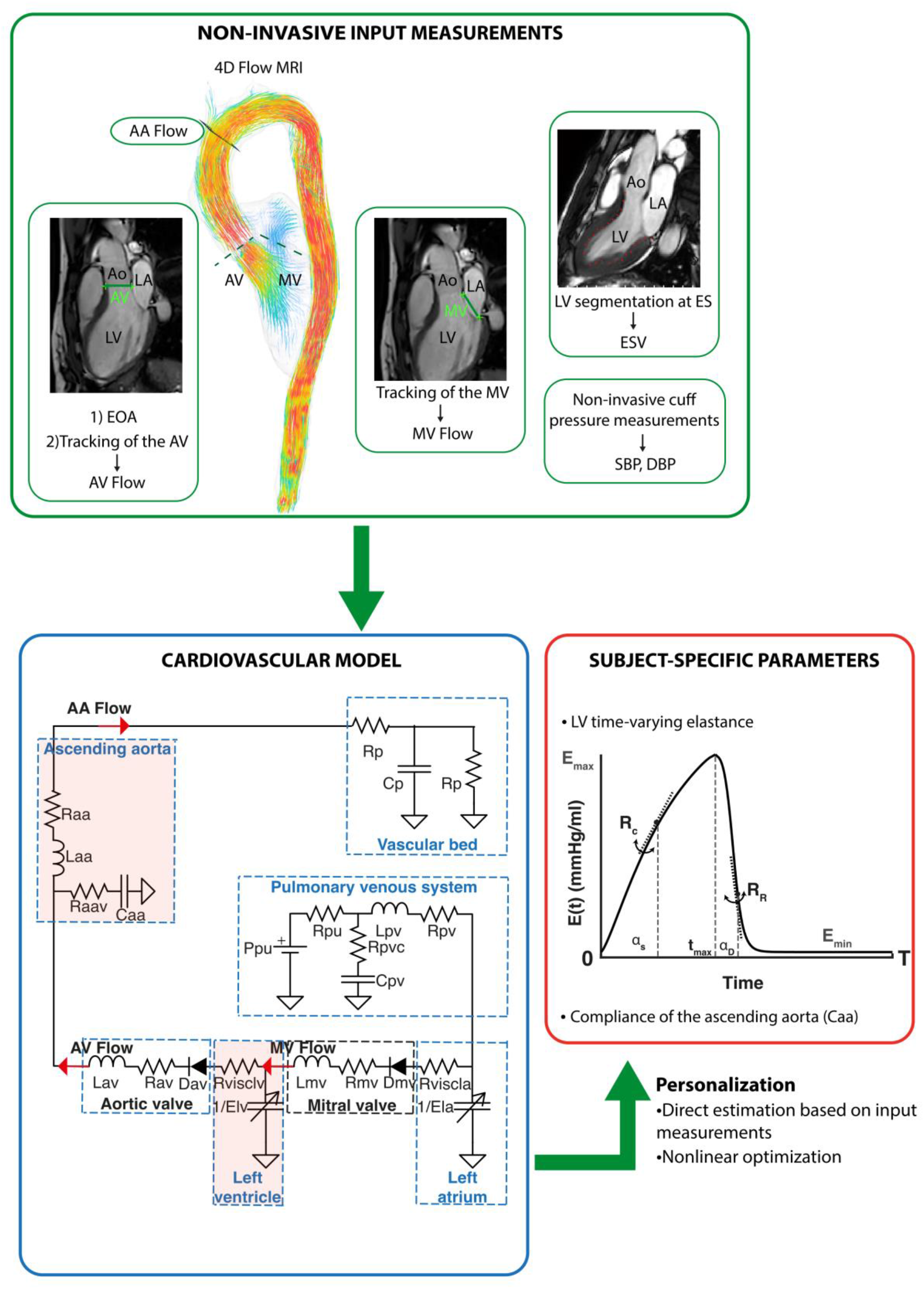
Schematic diagram of the personalization approach. The lumped parameter model is personalized using input measurements derived from the 4D Flow MRI data, the morphological CMR images and the non-invasive cuff pressure measurements in the brachial artery (SBP and DBP). Personalization yields subject-specific values of the model parameters, including the compliance of the ascending aorta (Caa) and parameters describing the LV time-varying elastance. The flow waveforms for personalization include the volumetric flows across the mitral valve (MV Flow), the aortic valve (AV Flow) and the ascending aorta, upstream from the brachiocephalic trunk (AA Flow). The time-varying elastance over the cardiac cycle length (T) is determined by the end-systolic, maximal elastance (*E*_*max*_), the minimal elastance (*E*_min_) over the duration of the cardiac cycle (T), the rates of LV contraction (*R*_*C*_) and relaxation (*R*_*R*_), and the systolic and diastolic time constants (*α*_*s*_ and *α*_*D*_, respectively). The maximal elastance occurs at the time of end-systole (*t*_max_). A description of the parameters in the model is given in Casas et al. ^6^. Ao, aorta; ES, End systole; EOA, effective orifice area; LA, left atrium; LV, left ventricle.

### Subject-specific parameter estimation

The model comprises 40 parameters, of which 23 were tuned to be subject-specific. The remaining 17 parameters were set to values reported in previous studies (Sun et al., 1995^3^; Heldt et al., 2002^4^; Garcia et al., 2012^38^; Mynard et al., 2012^39^; Broome et al., 2013^40^). The subject-specific parameters were obtained either by direct estimation from the input measurements or using a nonlinear optimization approach based on minimizing the error between the model-generated flow waveforms and those extracted from the 4D Flow MRI datasets. The number of parameters estimated using the nonlinear optimization routine was 20. Details of the optimization process are given in Casas et al.^6^.

### Intra- and inter-observer analysis and inter-sequence variability

The datasets were analyzed by two observers with four years of experience in cardiovascular MR. To evaluate the intra-observer variability, one observer (BC) performed two blinded analyses of the morphological and 4D Flow images for the ten study subjects. The analysis included computation of the ESV, the EOA, and the volumetric flow waveforms at the mitral valve, the aortic valve, and the ascending aorta. Inter-observer analysis was performed independently by a second observer (FV) who repeated the analysis on the same set of images. For the inter-scan variability, the second observer (FV) analyzed the 4D Flow MRI datasets acquired with the SGRE and the EPI sequences for the ten subjects in the study. This analysis involved calculation of the EOA and the flow waveforms at the mitral valve, the aortic valve, and the ascending aorta. The input data derived from the intra-observer, inter-observer, and inter-sequence analyses were used as input to estimate the subject-specific model parameters and evaluate their variability.

### Statistical analysis

Statistical analysis was performed using SPSS Statistics v.23.0 (IBM Corp., Armonk, NY, USA). All data are reported as mean ± standard deviation (SD) unless otherwise stated. Mann-Whitney U test for non-normally distributed data were performed for comparing image-derived measurements and model parameters obtained in the intra- and inter-observer studies, as well as from different sequences. A *P* value <0.05 was considered significant. The reproducibility of the results was evaluated using the coefficient of variation (CoV) ^41^. In addition, Bland-Altman plots were used to visually assess the agreement between the results. The bias ^*d̅*^ and the limits of agreement (^*d̅*^ ± 1.96*SD*) were derived from the Bland-Altman analysis.

## ETHICS STATEMENT

This study was carried out in accordance with the recommendations of the Regional Ethical Review Board in Linköping, with written informed consent from all subjects. All subjects gave written informed consent in accordance with the Declaration of Helsinki. The protocol was approved by the Regional Ethical Review Board in Linköping (Dnr 2015/396-31).

## ACKNOWLEDGEMENTS

This work was supported by grants from European Research Council; Grant number: 310612, the Swedish Research Council; Grant number: 2022-03931, Swedish Heart and Lung Foundation, 20210441 and 20230201, and ALF Grants Region Östergötland, number RÖ-987498.

## AUTHOR CONTRIBUTIONS

B.C, G.C, M.K and T.E participated in the conception and design of the study. B.C and F.V were involved in the analysis of the input data to the model. B.C performed the computational analysis and drafted the manuscript. B.C, F.V, K.T, G.C, C-J.C, MK and T.E interpreted the results. All authors edited and revised the manuscript. All authors read and approved the final version of the manuscript.

## DATA AVAILABILITY

The MRI datasets used to construct the models are available from the Linköping University Hospital for researchers who meet the criteria for access to confidential data. The codes used for simulating the model, as well as the optimization algorithms for personalization, are available from the author upon request.

