## Supplementary for "Reproducibility of 4D Flow MRI-based Personalized Cardiovascular Models; Inter-sequence, Intra-observer, and Inter-observer variability"

**SUPPLEMENTARY MATERIAL**

| | Descriptive Statistics | | Limits of agreement | |
|---|---|---|---|---|
| | ***Mean ± SD*** | $\bar{d}$ | ***-1.96*SD*** | ***+1.96*SD*** |
| **Intra-Observer Agreement** | | | | |
| *Left ventricle* | | | | |
| Passive LV elastance (mmHg/mL) | 0.09 ± 0.02 | -0.001 | -0.03 | 0.03 |
| Maximal LV elastance (mmHg/mL) | 2.2 ± 0.6 | -0.03 | -0.3 | 0.2 |
| Rate of ventricular contraction (-) | 1.5±0.3 | -0.03 | -0.3 | 0.3 |
| Rate of ventricular relaxation (-) | 30.3 ± 3.9 | -0.8 | -6.7 | 5 |
| Ventricular systolic time constant (-) | 0.4 ± 0.1 | -0.02 | -0.1 | 0.1 |
| Ventricular diastolic time constant (-) | 0.4 ± 0.03 | -0.006 | -0.04 | 0.03 |
| *Ascending aorta* | | | | |
| Compliance of the ascending aorta (mL/mmHg) | 0.1 ± 0.04 | -0.002 | -0.04 | 0.04 |
| **Inter-observer agreement** | | | | |
| *Left ventricle* | | | | |
| Passive LV elastance (mmHg/mL) | 0.09 ± 0.02 | -0.004 | -0.04 | 0.03 |
| Maximal LV elastance (mmHg/mL) | 2.2 ± 0.5 | -0.05 | -0.4 | 0.3 |
| Rate of ventricular contraction (-) | 1.5 ± 0.3 | -0.08 | -0.5 | 0.3 |
| Rate of ventricular relaxation (-) | 30.3 ± 3.9 | 0.2 | -5.3 | 5.7 |
| Ventricular systolic time constant (-) | 0.4 ± 0.1 | -0.04 | -0.2 | 0.2 |
| Ventricular diastolic time constant (-) | 0.4 ± 0.04 | -0.01 | -0.06 | 0.04 |
| *Ascending aorta* | | | | |
| Compliance of the ascending aorta (mL/mmHg) | 0.1± 0.04 | 0.003 | -0.07 | 0.07 |

**Table 1:** Bland-Altman analysis comparing the model-based parameters from the intra-and inter-observer study. The bias $\bar{d}$ and the limits of agreement ($\bar{d} \pm 1.96SD$) were derived from the analysis.

| | Descriptive Statistics | | Limits of agreement | |
|---|---|---|---|---|
| | ***Mean ± SD*** | $\bar{d}$ | ***-1.96*SD*** | ***+1.96*SD*** |
| **Inter-Sequence Agreement** | | | | |
| *Left ventricle* | | | | |
| Passive LV elastance (mmHg/mL) | 0.09 ± 0.03 | 0.003 | -0.04 | 0.05 |
| Rate of ventricular contraction (-) | 1.6±0.2 | -0.05 | -0.5 | 0.5 |
| Rate of ventricular relaxation (-) | 30.4 ± 4.4 | -0.5 | -6.8 | 5.9 |
| Ventricular systolic time constant (-) | 0.4 ± 0.1 | 0.003 | -0.2 | 0.2 |
| Ventricular diastolic time constant (-) | 0.4 ± 0.04 | -0.0001 | -0.05 | 0.05 |
| *Ascending aorta* | | | | |
| Compliance of the ascending aorta (mL/mmHg) | 0.1 ± 0.04 | 0.02 | -0.08 | 0.1 |

**Table 2:** Bland-Altman analysis comparing the model-based parameters from the inter-sequence study. The bias $\bar{d}$ and the limits of agreement ($\bar{d} \pm 1.96SD$) were derived from the analysis.
